# IPSE, a urogenital parasite-derived immunomodulatory protein, ameliorates ifosfamide-induced hemorrhagic cystitis through downregulation of pro-inflammatory pathways

**DOI:** 10.1101/381764

**Authors:** Evaristus C. Mbanefo, Loc Le, Rebecca Zee, Nirad Banskota, Kenji Ishida, Luke F. Pennington, Justin I. Odegaard, Theodore S. Jardetzky, Abdulaziz Alouffi, Franco H. Falcone, Michael H. Hsieh

## Abstract

Ifosfamide and other oxazaphosphorines can result in hemorrhagic cystitis, a constellation of complications caused by acrolein metabolites. We previously showed that a single dose of IPSE, a schistosome-derived host modulatory protein, can ameliorate ifosfamide-related cystitis; however, the exact mechanisms underlying this urotoxic effect and its prevention are not fully understood. To provide insights into IPSE’s protective mechanism, we undertook transcriptional profiling of bladders from ifosfamide-treated mice, with or without IPSE pretreatment. Following ifosfamide challenge, there was upregulation of a range of pro-inflammatory genes. The pro-inflammatory pathway involving the IL-1*β*, TNF*α* and IL-6 triad via NFκB and STAT3 signaling pathways was identified as the key driver of inflammation. The NRF2-mediated oxidative stress response pathway, which regulates both *Hmox1*-mediated heme homoeostasis and expression of antioxidant enzymes, was highly activated. Anti-inflammatory and cellular proliferation cascades implicated in tissue repair, namely Wnt, Hedgehog and PPAR pathways, were downregulated. IPSE administration before ifosfamide injection resulted in significant downregulation of major proinflammatory pathways including the triad of IL-1*β*, TNF*α* and IL-6 pathways, the interferon signaling pathway, and less apparent reduction in oxidative stress responses. Taken together, we have identified signatures of acute phase inflammation and oxidative stress responses in the ifosfamide-injured bladder, which are reversed by pretreatment with IPSE, a parasite derived anti-inflammatory molecule. In addition to providing new insights into the underlying mechanism of IPSE’s therapeutic effects, this work has revealed several pathways that could be therapeutically targeted to prevent and treat ifosfamide-induced hemorrhagic cystitis.

## Introduction

Hemorrhagic cystitis is a serious and difficult to manage complication resulting from exposure to certain chemotherapeutic agents (1-4), radiation therapy (4-7) and various viruses in immunosuppressed patients (8-11). Indeed, anticancer doses of oxazaphosphorines, such as cyclophosphamide and ifosfamide, are limited in part due to the risks of this complication (1-3). In the case of these agents, hepatic drug metabolism generates toxic acrolein which accumulates in bladder urine (12, 13). Fortunately, risks of chemotherapy-induced hemorrhagic cystitis have been decreased through the use of 2-mercaptoethane sulfonate Na (MESNA), which directly binds and neutralizes acrolein (14-16). However, MESNA fails to treat established hemorrhagic cystitis (14, 15, 17) and can also produce its own adverse reactions (18, 19). Other treatments options, including intravesically administered drugs (20-23), systemically administered agents (14, 24, 25), and nonpharmacological interventions (12, 24, 26-29), are either investigational or feature significant potential side effects (12, 30, 31).

The mechanisms underlying the initiation and pathogenesis of the acrolein-induced urotoxic effect are only partially elucidated. Knowledge gained from various studies and as reviewed by Haldar *et al.* (12) has implicated pro-inflammatory, heme homeostasis, and oxidative stress response pathways in the pathogenesis of acrolein-triggered bladder damage. Accumulation of acrolein-containing urine in the bladder lumen depletes the mucosal glycosaminoglycan layer and the asymmetric unit membrane (uroplakin complex), exposing the urothelium. Acrolein induces pyroptosis in the urothelium, a highly inflammatory form of apoptosis. The resulting sloughing and denudation of the urothelial layer exposes the lamina propria, detrusor muscle, and the bladder vasculature to further damage (12). Acrolein catalyzes reactions that generate reactive oxygen and nitrogen species (ROS and RNS) and superoxide radicals in the urothelium, resulting in membrane damage, DNA damage and cell death via the NFκB pathway (32, 33). The activation and involvement of the inflammasome complex in response to this oxidative stress results in the maturation and release of IL-1*β*, which in turn orchestrates a pro-inflammatory microenvironment in the urothelium (34, 35). This stress state also stimulates innate immune pattern recognition receptors (TLRs, NLRS and CLRs), sending signals that activate the NF*κ*B, STAT3, MAPK and other pro-inflammatory pathways, which lead to transcription of several pro-inflammatory cytokines (IL-1*β*, TNF*α* and IL-6), pro-inflammatory mediators (iNOS and COX-2) and chemokines that promote leukocyte infiltration and further drive inflammation and oxidative stress (34-36). In response to hemorrhage from damaged blood vessels and accumulating superoxide radicals, the heme homeostasis pathway and the oxidative stress response pathways are fully activated via NRF2-mediated mechanisms (37, 38). The clotting, edema and constriction of the bladder results in hyperalgesia.

There is a significant need for additional approaches to prevent and treat chemotherapy-induced hemorrhagic cystitis. Several analogs of cyclophosphamide that may enhance cytostatic efficacy while limiting urotoxicity have been explored, albeit with limited success (13, 39, 40). Candidate drugs targeting the inflammatory IL-1*β*, TNF*α* and IL-6 triad (35, 41) and/or promoting oxidative stress responses show promise for ameliorating hemorrhagic cystitis but have not progressed beyond preclinical testing. Most efforts have been focused towards finding alternatives to MESNA, including anti-inflammatory molecules (42-49), hemostatic agents (50-52), antioxidants (37, 48, 49, 53-59), analgesics (60), anti-depressants (61), vasodilator (62), cytokines (25, 63, 64), platelet rich plasma (65, 66), nutritional approaches (67, 68), and plant extracts (45, 46, 48, 56, 68-71). These early-stage drug candidates target pro-inflammatory pathways, heme homeostasis pathway and anti-oxidant homoeostasis.

Another potential approach to treat chemotherapy-induced hemorrhagic cystitis is to administer IL-4 (25), a potent anti-inflammatory cytokine known to antagonize the IL-1*β*, TNF*α*, and IL-6 pathways. This finding led us to test and verify that a single dose of an IL-4-inducing, parasite-derived anti-inflammatory molecule (IPSE, the IL-4-inducing principle from *Schistosoma mansoni* eggs) ameliorated the inflammation, hemorrhage, and urothelial sloughing associated with ifosfamide-induced hemorrhagic cystitis (42). IPSE binds immunoglobulins, notably IgE on the surface of basophils and mast cells, inducing secretion of preformed IL-4 (72-74). However, we suspect IPSE may have additional mechanisms underpinning its ability to alleviate ifosfamide-induced hemorrhagic cystitis. IPSE also sequesters chemokines (75), which likely orchestrate anti-inflammatory responses. As an infiltrin possessing a nuclear localization sequence (NLS), IPSE is able to translocate into host cell nuclei to modulate host gene transcription (76-78). Given that the transcriptional changes during ifosfamide-induced hemorrhagic cystitis are largely unknown, and because the underlying mechanisms of IPSE’s protective effects remain to be elucidated, we undertook transcriptome-wide profiling of the bladder of ifosfamide-treated mice using RNA-Seq. Furthermore, we studied the gene expression dynamics in IPSE pretreated mice challenged with ifosfamide. Here, we show that key pro-inflammatory, heme homeostatic and oxidative stress response pathways are highly activated in the bladder following ifosfamide insult. Finally, we show that IPSE downregulates pro-inflammatory responses as a potential protective mechanism, in addition to its involvement in promoting urothelial repair.

## Results

### Ifosfamide-induced hemorrhagic cystitis is ameliorated by IPSE

We recently showed that a single dose of IPSE was comparable to administration of recombinant IL-4 or three doses of MESNA in alleviating ifosfamide-induced hemorrhagic cystitis (42). We used these established methods to obtain bladder samples for transcriptional profiling. Mice were administered: 1) saline or 2) IPSE, 24 hours before ifosfamide challenge, or 3) saline vehicle alone. Twelve hours following ifosfamide insult, bladder histopathology was analyzed in a blinded fashion. Compared to bladders from saline-treated mice (Fig. 1A), bladders from mice challenged with ifosfamide showed marked edema, dysregulated contraction, hemorrhage, and urothelial sloughing (Fig. 1B and D). Conversely, bladders from mice treated with IPSE before ifosfamide challenge were significantly protected from urothelial denudation and inflammation (Fig. 1C). Based on blinded scoring of bladder sections, we observed significant increases in inflammation (Fig. 1E), urothelial denudation (Fig. 1F), and edema (Fig. 1G), and non-statistically significant increases in hemorrhage (Fig. 1H) in ifosfamide-treated mice. These features were markedly reduced in mice administered a single dose of IPSE before ifosfamide treatment, in comparison to ifosfamide-treated mice. Both inflammation and urothelial denudation were significantly reduced (Fig. 1E and F), while edema and hemorrhage were reduced but not statistically significant (Fig. 1G and H). Taken together, this qualitative and quantitative data demonstrate characteristic features of ifosfamide-induced hemorrhagic cystitis, some of which were significantly reduced by IPSE pretreatment.

**Fig. 1.**
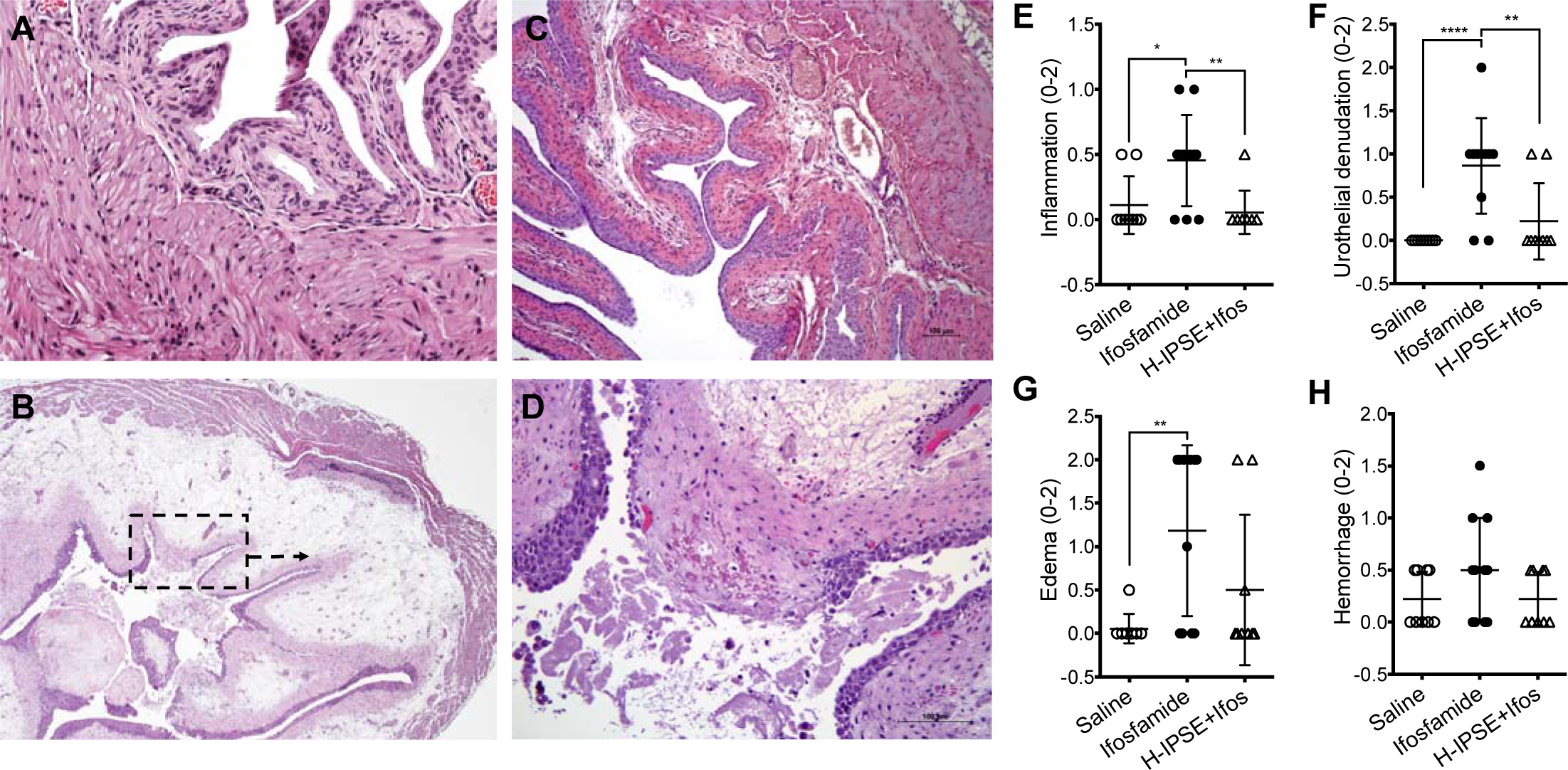
IPSE ameliorates ifosfamide-induced hemorrhagic cystitis. Mice were pretreated with saline or IPSE 24 hours before challenge with 400mg/kg of ifosfamide. Bladders were assessed for histopathologic changes following ifosfamide insult in a blinded fashion. (A) Normal bladder showing intact urothelium with no signs of pathology. (B) Bladder from an ifosfamide-treated mouse (pretreated with saline) showing urothelial sloughing and edema. (C) Bladder from an IPSE-pretreated, ifosfamide-challenged mouse showing significant reduction in inflammation, urothelial denudation and edema. (D) High power view of bladder section shown in dotted box in (B). Graphs showing treatment group differences in bladder (E) inflammation, (F) urothelial denudation, (G) edema, and (H) hemorrhage. Each symbol represents the score for an individual mouse. Cross bar for each group denotes mean score. * p<0.05, ** p<0.01, **** p<0.001 based on post-hoc Students t-tests following significant difference among groups by ANOVA.

### Transcriptional profiles show massive pro-inflammatory response and activation of oxidative stress responses during ifosfamide-induced hemorrhagic cystitis

Ifosfamide is metabolized in the liver to generate acrolein, which is secreted in urine and damages the bladder (18). To understand the transcriptional alterations elicited by acrolein in the bladder during ifosfamide-induced hemorrhagic cystitis, mice were treated with saline vehicle or ifosfamide. Gene expression dynamics in the ifosfamide-injured bladder were studied through RNA-Seq performed on bladders harvested 6 hours following ifosfamide injection. RNA sequencing was performed to a considerable depth (20 million reads), more than 96% of which were successfully aligned to the *Mus musculus* genome. Principal component analysis indicated gene expression homogeneity among ifosfamide-treated bladders relative to the vehicle control (Fig. 2A) and a slight overlap between bladders from ifosfamide-treated mice and IPSE pretreated mice challenged with ifosfamide (Fig. 2B). Volcano plotting of differentially expressed genes and their associated statistical significance (Fig. 2C) revealed upregulation of a large set of genes (n = 2061) and downregulation of an appreciable number of genes (n = 1114), based on *p-value* (adjusted) < 0.1 and *log2*(Fold Change) > 1 (at least 2-fold). Among the top upregulated genes were *Il6*, a major member of the IL-1*β*, TNF*α* and IL-6 pro-inflammatory triad. These three cytokines together are major drivers of inflammatory responses (see relationship with other downstream proinflammatory genes, regulators and mediators in Fig. 2D). Indeed, the *Il1b, Tnfa* and *Il6* genes were upregulated by about two orders of magnitude. The *Ptx3* gene was also one of the top upregulated genes; a member of the pentraxin protein family, major components of the humoral arm of innate immune response highly induced in response to inflammatory stimuli (79). Chemokines were also highly upregulated, especially *Cxcl2* and *Ccl2*. In addition, the *Hmox1* gene encoding the heme oxygenase 1 enzyme, the first enzyme of the heme oxygenase pathway, was also highly upregulated. The *Eid3* gene, also among the most upregulated genes, is involved in cellular responses to stress (Fig. 2C and Supplementary Fig. S1). The cysteine transporter, *Slc7a11,* which has been implicated in glutathione metabolism in the bladder (80, 81), was also significantly upregulated in ifosfamide-injured bladders (Fig. 2C).

**Fig. 2.**
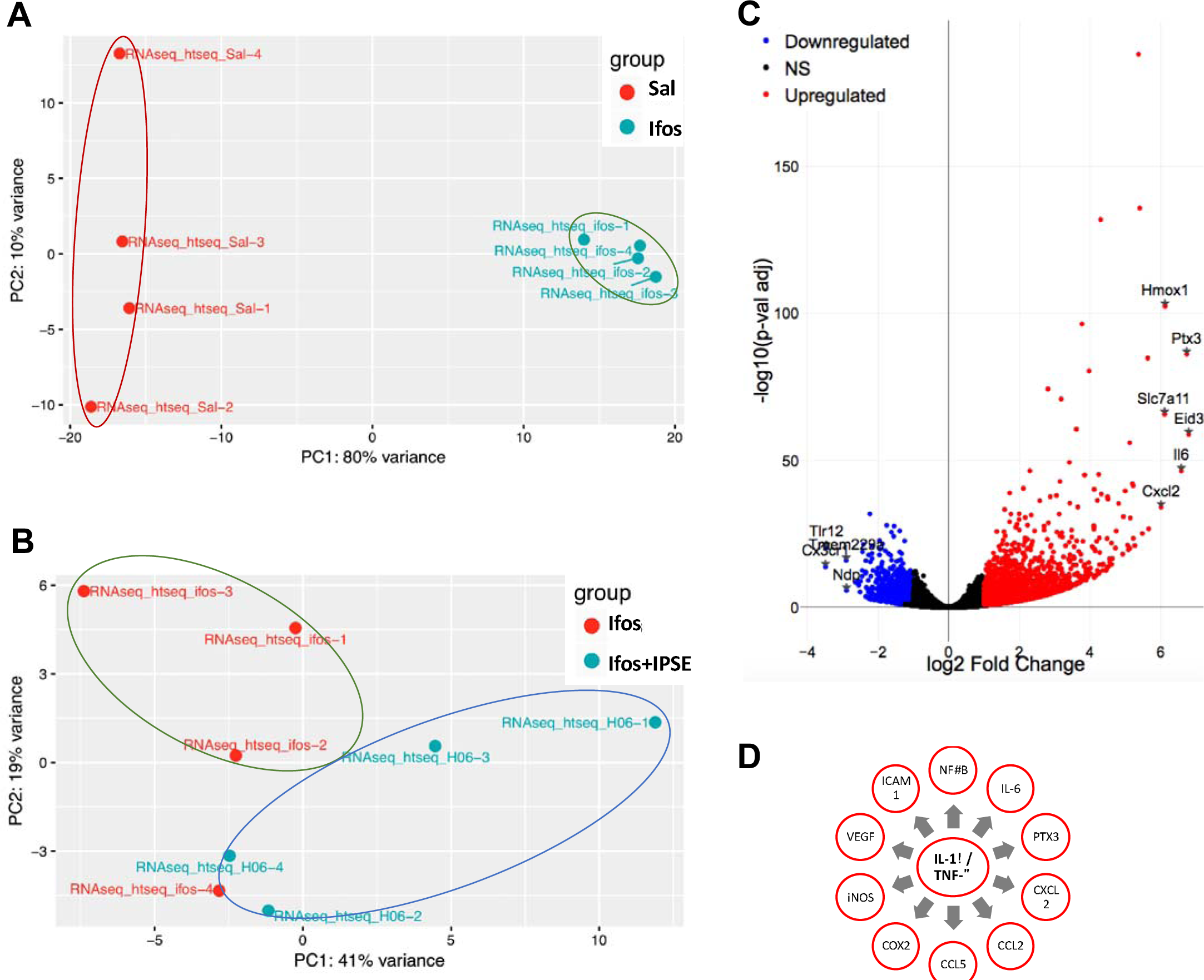
RNA-Seq analysis of ifosfamide-exposed bladders indicates multiple inflammation and stress response-related genes are differentially regulated. Transcriptional changes in bladders from mice administered saline were compared to those occurring in bladders from mice given ifosfamide. (A) Principal component analysis (PCA) showed homogeneous clustering of gene expression among ifosfamide-treated mice (turquoise symbols labeled with “ifos” suffix) and vehicle-treated mice (red symbols labeled with “Sal” suffix). (B) Principal component analysis (PCA) showed overlap of gene expression among ifosfamide-treated mice mice (red symbols labeled with “Ifos” suffix) and IPSE-treated mice challenged with ifosfamide (turquoise symbols labeled with “H06” suffix). (C) Volcano plots demonstrated upregulated and downregulated genes in bladders from ifosfamide-versus saline-treated mice. For this comparison, p-value (adjusted) < 0.1 and log2(Fold Change) > 1 (at least 2-fold) were applied as threshold values. NS: genes with non-significant changes in expression levels [black dots]. (D) Schematic representation of the relationships among the IL-1*β*, TNF*α* and IL-6 triad and downstream pro-inflammatory cytokines and mediators.

Pathway and functional analysis expectedly revealed signatures of inflammation. Specifically, there was differential activity involving the IL-6 pathway, in which IL-1*β*, TNF*α* and IL-6 play major roles, and other pro-inflammatory pathways implicated in the pathogenesis of ifosfamide-induced hemorrhagic cystitis (35, 41, 47, 82-84) (Fig. 2C, Fig. 3 and Supplementary Fig. S1). IL-6 and its cognate receptors were highly upregulated, in addition to STAT3 and the tyrosine protein kinase JAK2, which are both involved in IL-6 signaling (Fig. 4A and Supplementary Fig. S2A). Similarly, IL-1*β*, TNF*α* and their receptors were upregulated. These cascades converge through TAK1 to promote formation of the IκB-NFκB complex and drive pro-inflammatory gene transcription in conjunction with NF-IL-6, the nuclear factor of IL-6 expression (Fig. 4A and Supplementary Fig. S2B). Accordingly, the STAT3 and NFκB pathways, both major drivers of inflammation and immune response via the IL-1*β*-TNF*α*-IL-6 triad, were upregulated following ifosfamide insult transcription (Supplementary Fig. S2A and B). Other major upregulated pro-inflammatory pathways and disease signaling cascades included those related to TNF receptor, iNOS, the acute phase response, diabetes mellitus, HMGB1, oncostatin, generation of ROS and RNS, and major innate immune-related cascades (Fig. 3). Finally, there was noteworthy upregulation of the IL-17F-mediated allergic inflammatory and leukocyte extravasation signaling pathways (Supplementary Fig. S3).

**Fig. 3.**
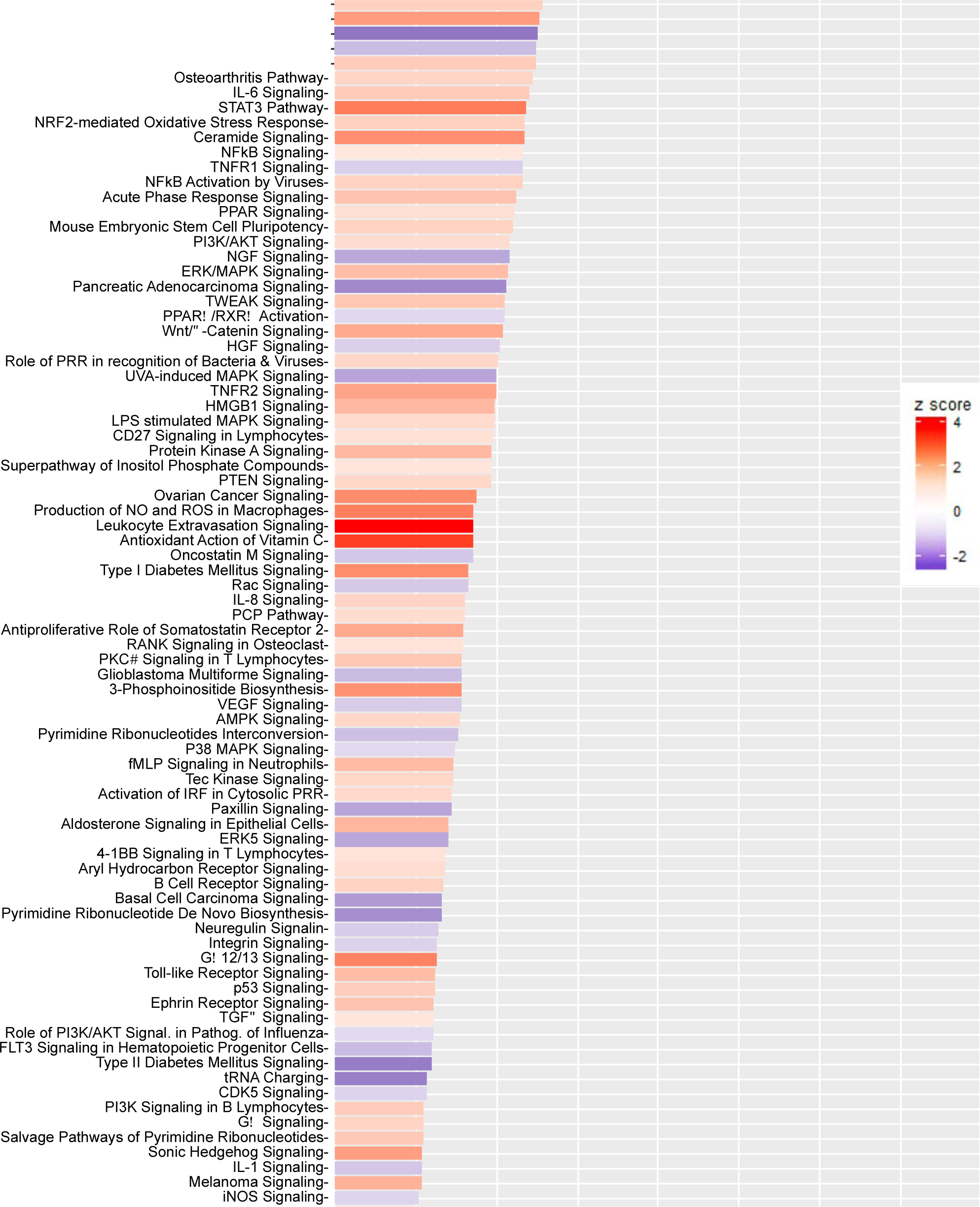
Most differentially altered gene pathways in the bladder during ifosfamide-induced hemorrhagic cystitis. Functional comparison of the transcriptome of bladders from ifosfamide-versus vehicle-treated mice was performed using Ingenuity Pathway Analysis. Bars are colored according to z-score, with red showing upregulation and blue denoting downregulation. The size of each bar is proportional to its –log(p-value).

**Fig. 4.**
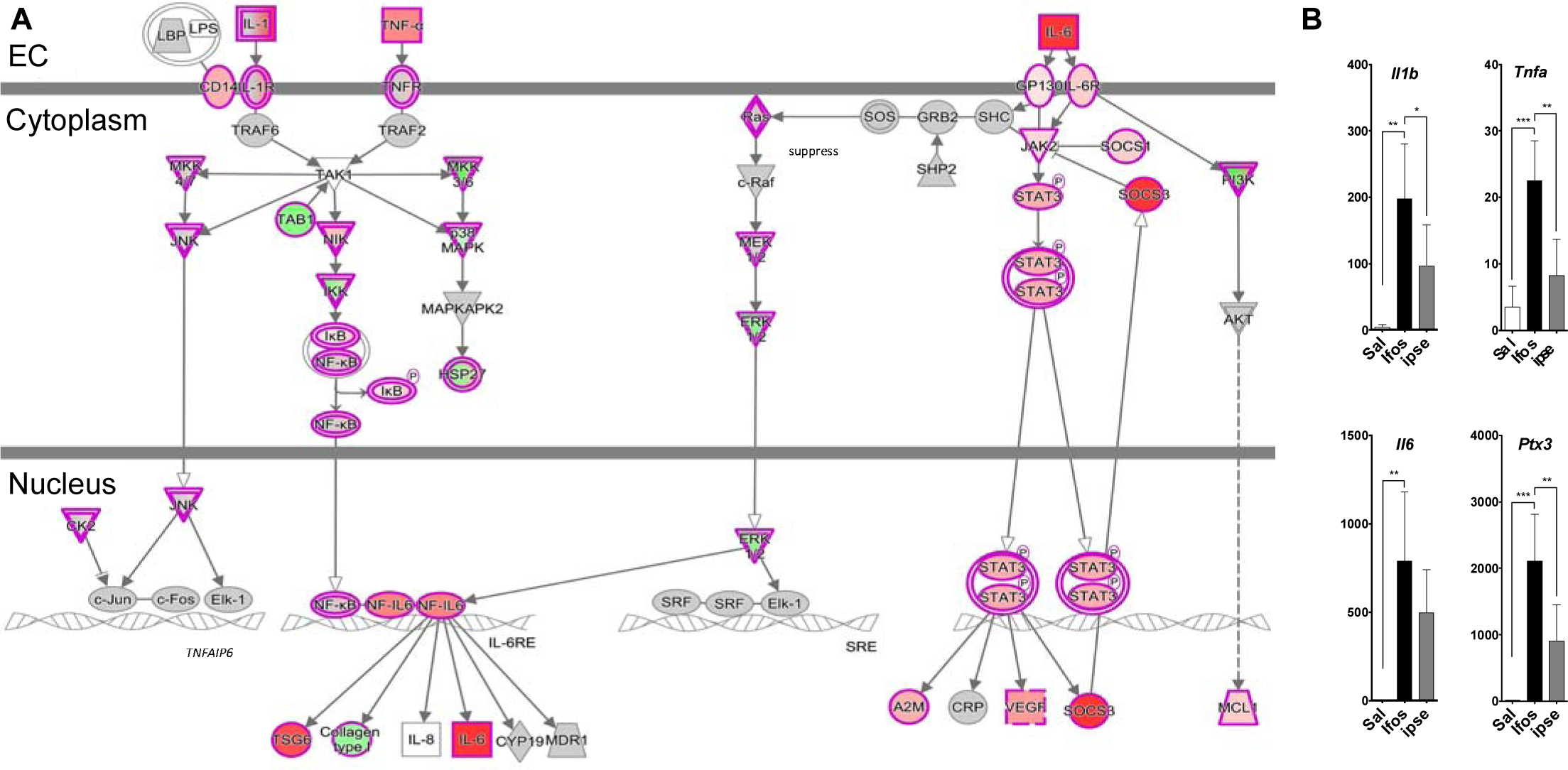
The IL-1*β*, TNF*α*, and IL-6 triad of pathways are major inflammatory gene pathways upregulated in the bladder during ifosfamide-induced hemorrhagic cystitis. (A) Bladders of ifosfamide-exposed mice upregulated expression of genes from the IL-1*β*, TNF*α* and IL-6 triad of pathways and their corresponding cytokines, receptors and downstream nuclear transcriptional factors. In the case of IL-1*β* and TNF*α*, these cascades converge upon NFκB. IL-6 also indirectly interacts with NFκB through ERK1/2 activation of NK-IL6, which works with NFκB to promote transcription of target genes. Keys: upregulation (red), downregulation (green), cytokines (square), growth factors (dotted square), phosphatase (triangle), kinases (inverted triangle), transmembrane receptors (ellipse), transcriptional regulators (wide circle), peptidase (rhombus), group or complex (double lined shapes), transporter (trapezium), acts on (line with filled arrow), translocate (line with open arrow), inhibition (line with perpendicular line at edge). (B) Both IL-1*β* and TNF*α* gene transcription were increased by ~100 fold in the bladders of ifosfamide-treated mice. Pretreatment with IPSE reduced the level by ~50% relative to the ifosfamide-treated group. Similar trends were observed for cognate receptors and downstream transcription factors (data not shown).

Another hallmark of ifosfamide-induced hemorrhagic cystitis is a significant oxidative stress response to acrolein exposure and resulting hemorrhage (33). Our data underscores a major role for the erythroid-derived leucine zipper NRF2 as a nuclear factor involved in regulating oxidative stress responses to ifosfamide injury of the bladder (37, 51, 53) (Fig. 3, 5A and Supplementary Fig. S4). The NRF2-mediated oxidative stress response gene pathway, which regulates the expression of antioxidant and heme homeostatic proteins, was one of the most upregulated pathways in the bladder following ifosfamide insult (Fig. 3, 5A and Supplementary Fig. S4). We noted considerable upregulation of genes encoding enzymes including heme oxygenase (HO-1), which catalyzes the first step in heme homeostasis, the antioxidant thioredoxin reductase (TRXR1), which catalyzes the reduction of thioredoxin to restore redox homeostasis, peroxiredoxin (PRDX1), which detoxifies peroxide radicals, and glutathione reductase (GSR), which reduces glutathione disulfide to glutathione, an important antioxidant that scavenges hydroxyl radicals. Also upregulated were thioredoxin (TXN), superoxide dismutase (SOD) and ferritin light and heavy chain proteins (FTL and FTH1), which are involved in redox signaling, superoxide partitioning and iron homeostasis, respectively (Fig. 5A). Genes encoding proteins involved in xenobiotic detoxification were also upregulated in response to acrolein (Fig. 5A). In addition, the p38 MAPK pathway, implicated in responses to stress stimuli (43), was also substantially upregulated.

**Fig. 5.**
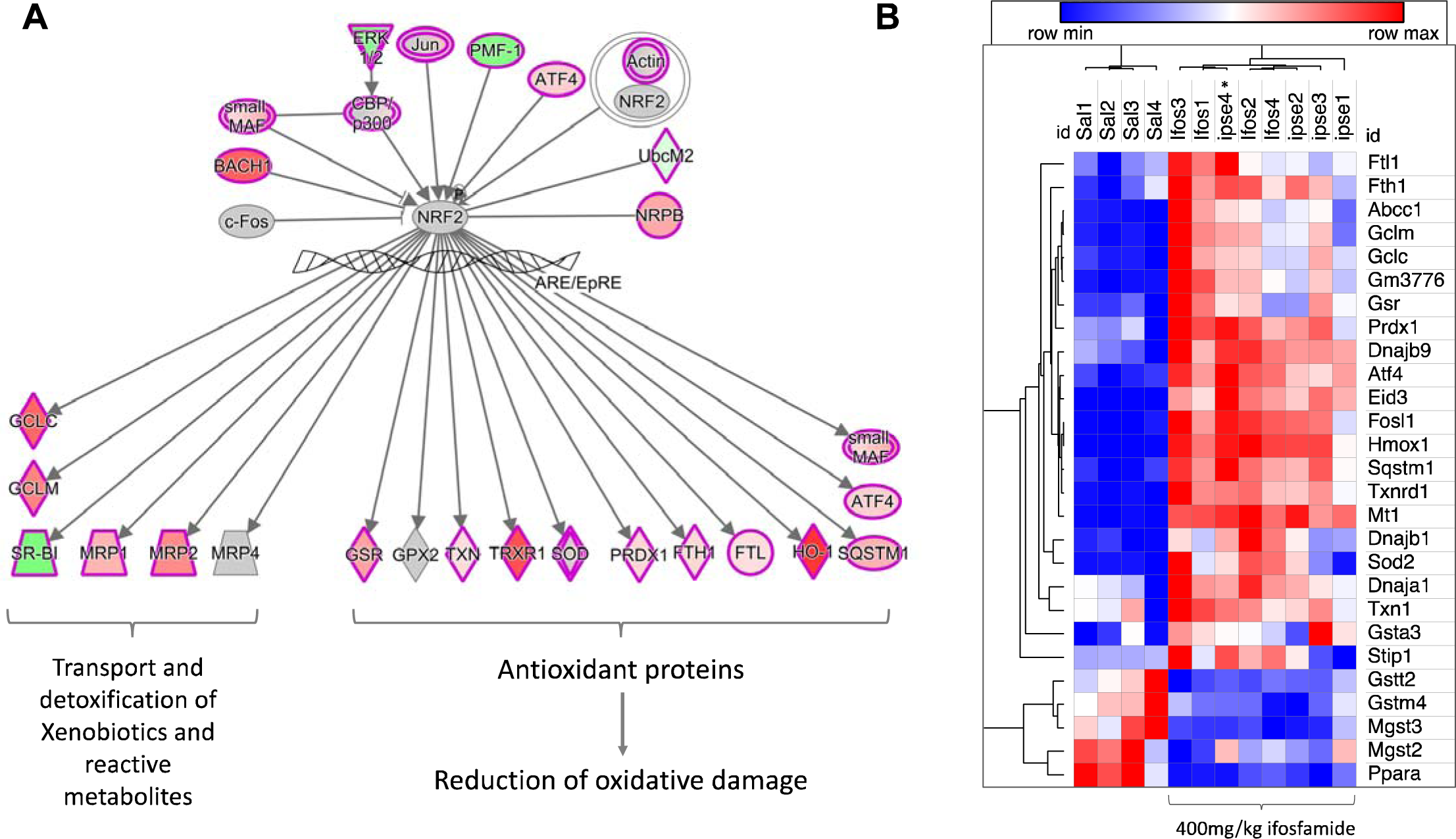
Oxidative stress responses of the bladder during ifosfamide-induced hemorrhagic cystitis. (A) Schematic representation of the relationships between NRF2 and antioxidant proteins and proteins involved in heme homeostasis and xenobiotic detoxification. A more detailed version is shown in Supplementary Fig. S4. The keys to the shapes and colors are as detailed in Fig. 4. (B) Heat map showing levels of expression of genes encoding major antioxidant enzymes. There were no overt changes in the levels of expression of these genes, except for Fth1, Ftl1, Abcc1, Gclc, Gclm, Gm3776 and to some extent for Gsr, Sod2 and Dnajb1. Red designates gene upregulation while blue denotes downregulation. The columns represent data for individual mice in each treatment group (“Sal”: saline vehicle treatment, “Ifos”: ifosfamide treatment only, “Ipse”: ifosfamide and IPSE treatment).

The most downregulated pathway in the ifosfamide-exposed bladder was the peroxisome proliferator-activated receptor (PPAR) signaling pathway, which is involved in lipid homoeostasis (85), in addition to its anti-inflammatory effect (86, 87) and role in the development and maintenance of IL-4 dependent alternatively activated status in macrophages (88) (Fig. 3 and Supplementary Fig. S5). TWEAK, Wnt and Hedgehog, which can mediate anti-inflammatory responses, were likewise downregulated (Fig. 3). The aldosterone signaling in epithelial cells pathway, which is involved in ion transport to maintain electrolyte and water balance across epithelial surfaces, was also downregulated (Fig. 3). Finally, analysis of diseases and functions affected by bladder ifosfamide challenge showed considerable upregulation of functions related to organismal injury and abnormalities, inflammatory diseases, cancer, cell proliferation, cellular movement and hematological development and function (Supplementary Fig. S6). There was also notable upregulation of genes and regulators in the neuro-inflammatory pathways (Supplementary Fig. S7). Analysis of differentially expressed gene members of neuro-inflammatory pathways indicated significant expression of genes encoding proinflammatory cytokines and mediators in neuronal cells. Although this may suggest role in the acrolein-associated hyperalgesia, the presence of astrocytes or microglia in the peripheral nervous system within the bladder is unproven (Supplementary Fig. S7). Finally, the HIF-1*α*-mediated hypoxia-related signaling cascade, was upregulated in the bladder after ifosfamide insult, consistent with the hemorrhage associated with hemorrhagic cystitis (Supplementary Fig. S8).

### Transcriptional modulatory effects of IPSE on the ifosfamide-exposed bladder

We recently reported that IPSE, an immunomodulatory protein of parasite origin, can ameliorate much of the pathology associated with ifosfamide-induced hemorrhagic cystitis ((42) and Fig. 1). To provide insight into the underlying mechanisms of the IPSE’s protective effects, we undertook gene expression profiling of the ifosfamide-challenged bladder, with or without IPSE pretreatment. Mice were treated with saline or IPSE, 24 hours before challenge with ifosfamide. Gene expression dynamics were profiled through RNA-Seq analysis of bladders harvested 6 hours following ifosfamide administration. Compared to mice receiving ifosfamide without IPSE pretreatment, genes encoding cytokines driving pro-inflammatory responses (IL-1*β*, TNF*α* and IL-6 triad) were downregulated 50% in the bladders of mice treated with IPSE before ifosfamide challenge (Fig. 4B). Similar downward trends in gene expression were observed for these cytokine’s receptors and downstream genes and other genes and transcriptional factors driving inflammation (Fig. 4B and 6A). The expression of *Cxcl10* (IP-10), a major interferon gamma-induced chemokine, was highly increased in ifosfamide-injured bladders but downregulated in the bladders of IPSE-pretreated, ifosfamide-exposed mice (Fig. 6A). The transcriptional levels of several other chemokines also involved in inflammatory responses and recruitment of cells to sites of stress were highly upregulated in bladders only exposed to ifosfamide, but conversely downregulated in ifosfamide-injured bladders pretreated with IPSE (Fig. 6A). The predicted gene interaction grids associated with IPSE administration featured a network linking downregulation of *Ccl2* (and other chemokines genes) to the downregulation of a number of gamma interferon-inducible proteins and nitric oxide synthase (Fig. 6B and Supplementary Fig. S9). Other mechanistic networks suggesting IPSE induced downregulation of additional pro-inflammatory factors were also noted (Supplementary Fig. S9).

**Fig. 6.**
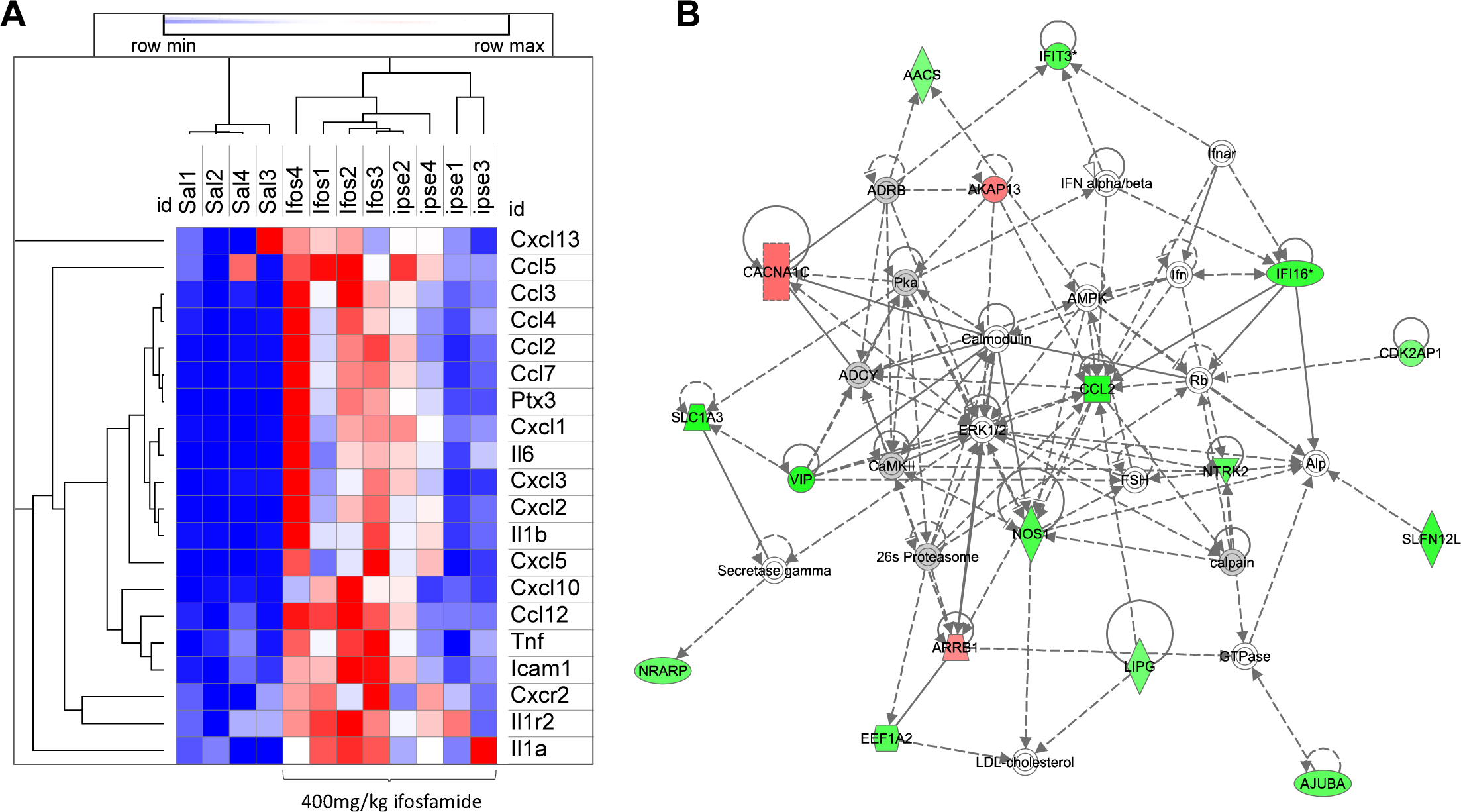
Effects of IPSE on bladder transcription of inflammation-related genes during ifosfamide-induced hemorrhagic cystitis. (A) Heat map showing levels of expression of genes encoding major proinflammatory proteins, their receptors, and downstream proinflammatory mediators and chemokines. IPSE pretreatment resulted in significant downregulation of major proinflammatory proteins, their receptors, and downstream proinflammatory mediators and chemokines. Red designates gene upregulation while blue denotes downregulation. The columns represent data for individual mice in each treatment group (“Sal”: saline vehicle treatment, “Ifos”: ifosfamide treatment only, “Ipse”: ifosfamide and IPSE treatment). (B) A representative mechanistic network showing inhibitory relationships among chemokines (Ccl2), nitric oxide synthase and several interferon-induced proteins. The keys to the shapes and colors are as detailed in Fig. 4.

The most downregulated gene pathway (in terms of statistical significance) in the bladders of IPSE-pretreated, ifosfamide-exposed mice, compared to bladders exposed only to ifosfamide, was the interferon signaling pathway (Fig. 7). Many of the major gene pathways noted to be highly upregulated in ifosfamide-damaged bladders were relatively downregulated in bladders pretreated with IPSE before ifosfamide challenge (Fig. 3 and 7). These downregulated pathways included those related to interferon signaling, inflammatory diseases such as osteoarthritis and diabetes mellitus, pattern recognition receptor signaling pathways of the innate immune system, pro-inflammatory pathways including NFκB, iNOS, neuro-inflammation, TREM1, the acute phase response, HMGB1, STAT3, IL-6, TNFR, IL-1 and pathways involved in the production of ROS and RNS (Fig. 7). We also observed a relative increase in metabolic gene expression relevant to oxidative phosphorylation, glycolysis and PPAR signaling. Interestingly, the most downregulated genes in terms of z-score were those related to the neuro-inflammation signaling pathway (astrocytes and microglia). While a direct effect of IPSE on bladder neurons could not be inferred based on this finding, due to a lack of astrocytes and microglia in the bladder, we have observed significant reduction in bladder pain in IPSE-treated mice challenged with ifosfamide (42).

**Fig. 7.**
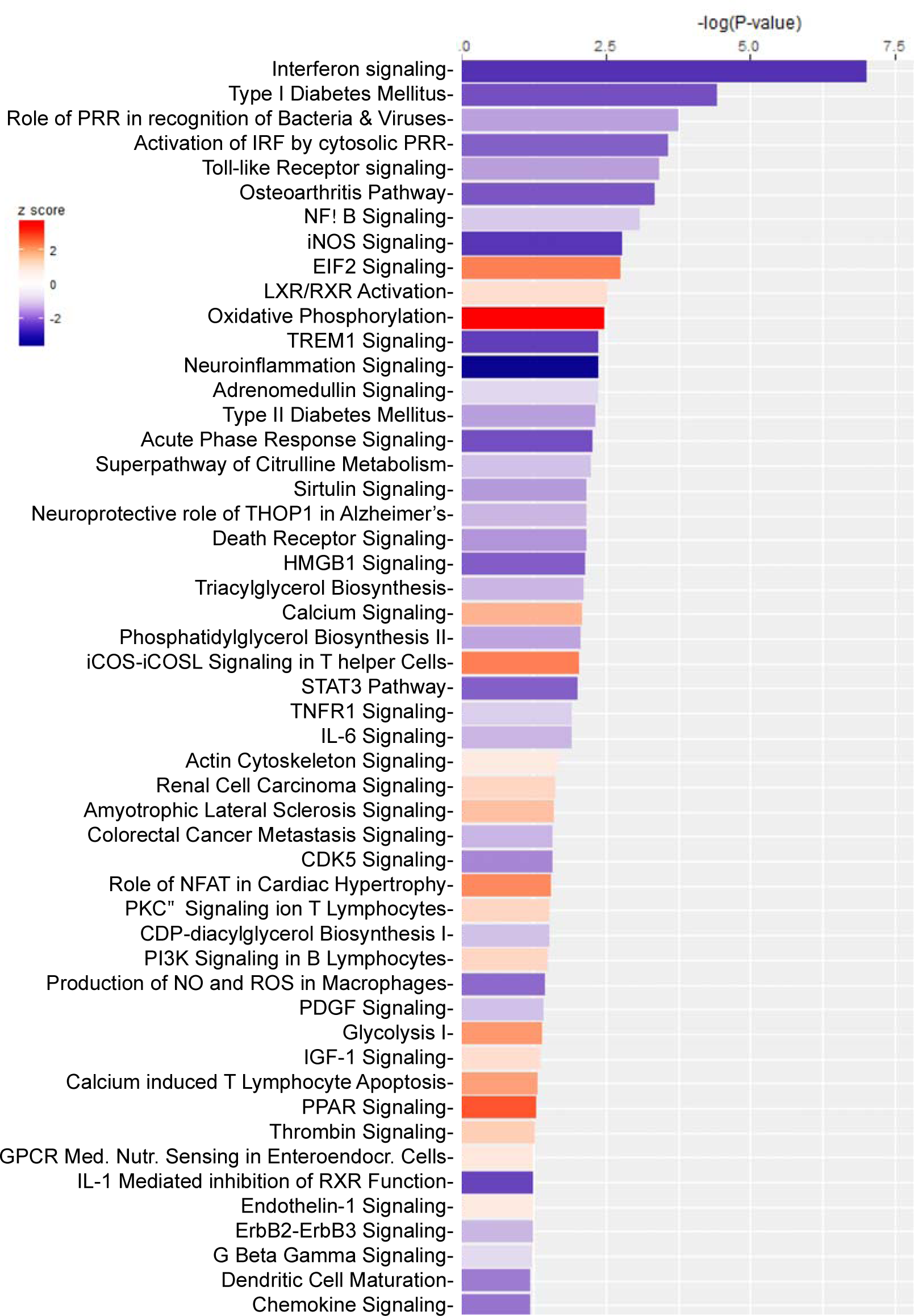
Most differentially altered gene expression pathways in bladders from IPSE-pretreated mice challenged with ifosfamide. Mice were pretreated with saline or IPSE, 24 hours before challenge with 400mg/kg of ifosfamide. The bladders were subjected to transcriptional profiling (RNA-Seq) and functional analysis using Ingenuity Pathway Analysis. Bars are colored according to z-score, with red showing upregulation and blue denoting downregulation. The size of each bar is proportional to its –log(p-value).

Compared to bladders pretreated with IPSE before ifosfamide insult, bladders exposed only to ifosfamide relatively upregulated gene expression of some antioxidant enzymes, including *Fth1, Ftl1, Abcc1, Gclc, Gclm, Gm3776* and to some extent for *Gsr, Sod2* and *Dnajb1* (Fig. 5B). It is notable that the downregulated gene expression of proteins in this pathway were those involved in DNA damage sensing, superoxide partitioning and detoxification of xenobiotics, and metal ion homeostasis (Fig. 5B). In particular, relatively lower expression of genes encoding the ferritin proteins (*Fth1* and *Ftl1)* suggest relatively earlier restoration of iron homeostasis in IPSE pretreated mice, supporting the observed decrease in bladder hemorrhage induced by IPSE pretreatment before IFS challenge (42). Taken together, these data suggest in IPSE-pretreated bladders exposed to ifosfamide, that there is a significant anti-oxidant response following accumulation of acrolein, but lower expression of genes related to detoxification of xenobiotics, DNA damage sensing and iron homeostasis.

## Discussion

Herein we describe the first transcriptome-wide profiling of the bladder during ifosfamide-induced hemorrhagic cystitis. To accomplish this, we used a tractable mouse model which recapitulates the pathogenesis of hemorrhagic cystitis (25) resulting from the urotoxic effect of acrolein, a byproduct of ifosfamide metabolism. This study has verified a number of important findings regarding specific biological aspects of ifosfamide-induced hemorrhagic cystitis. We have expanded upon this body of work by defining multiple key pathogenetic mechanisms through comprehensive transcriptomics. Furthermore, our RNA-Seq data extends our prior work on the therapeutic effect of IPSE in ifosfamide-induced hemorrhagic cystitis (42). This study has revealed a central role played by the IL-1*β*, TNF*α* and IL-6 triad in driving the substantial inflammation associated with ifosfamide-induced hemorrhagic cystitis (35, 41, 47, 82-84). A 100-fold increase in expression of IL-1*β*, TNF*α* and Il-L-6 was reduced ~50% by pretreatment with a single dose of IPSE. We also confirmed that gene members of the heme hemostasis and oxidative stress response biological systems were highly transcribed following ifosfamide-induced hemorrhagic cystitis, presumably to restore antioxidants to normal levels (37, 48, 50-53, 56-58). Moreover, we have shown that these urotoxicity-associated transcriptional changes, especially inflammatory responses, were downregulated by IPSE when administered before ifosfamide challenge.

Unlike MESNA, which binds to and neutralizes acrolein directly to prevent urotoxicity, IPSE prevents or reverses the inflammatory changes that drive bladder damage following ifosfamide exposure. We postulate that IPSE, through its inhibitory transcriptional effects on IL-1*β*, TNF*α* and IL-6, key upstream cytokines driving inflammation via NFκB and STAT3, limits ifosfamide-triggered inflammation, urothelial denudation and vascular pathogenesis. The ability of IPSE to induce gene expression of uroplakins (42), crucial urothelial barrier function genes, is a likely contributor to IPSE’s therapeutic effect on the bladder following chemical insult (59, 89, 90). Other candidate drugs for ifosfamide-induced hemorrhagic cystitis have also been shown to specifically target the IL-1*β*-TNF*α*-IL-6 pathway. Dantas *et al*. (2010) showed that simvastatin attenuated cyclophosphamide-induced urothelial inflammation by decreasing the expression and activities of IL-1*β*, TNF*α* and IL-6 (47). Recombinant IL-4, quinovic acid glycosides, oleuropein, anakinra, pentoxifylline, diallyl disulfide and other anti-inflammatory candidates were separately shown to reduce the pathogenesis of hemorrhagic cystitis by inhibition of the expression of these inflammatory cytokines and their receptors (25, 41, 43, 45, 48, 49). In accord with our observations, other drug studies have identified a therapeutic requirement for downregulation of IL-1*β*, TNF*α* and IL-6 associated transcriptional factors (NFκB and STAT3) (32, 43) and downstream inflammatory mediators (iNOS and COX-2) (25, 32, 48, 83, 84, 91, 92).

Besides effects on the IL-1*β*, TNF*α* and IL-6 triad, IPSE may also mediate critical gene expression changes in chemokines in ifosfamide-induced hemorrhagic cystitis. A range of major chemokines genes (*Ccl2*, *Ccl3, Ccl4, Ccl5, Ccl7, Ccl12, Cxcl1, Cxcl2, Cxcl3, Cxcl5, Cxcl10 and Cxcl13)* acting as chemo-attractants to site of stress were significantly downregulated by IPSE pretreatment before ifosfamide insult. For instance, *Cxcl10* transcription is upregulated in the bladder following ifosfamide insult, and is significantly decreased by comparison in IPSE-pretreated, ifosfamide-exposed bladders. Indeed, CXCL10 blockade has been shown to significantly dampen cyclophosphamide-induced hemorrhagic cystitis (93). Interleukin 8 receptor (*Cxcr2)* was also relatively downregulated by IPSE pretreatment before ifosfamide challenge. Notably, *Cxcr2* was previously identified to play an important role in cyclophosphamide induced hemorrhagic cystitis (94).

IPSE likely orchestrates a portion of its therapeutic effects through actions on other cytokines. Gene expression network analysis revealed a link between the observed downregulation of CCL2 in the bladders of IPSE-pretreated, ifosfamide-challenged mice (versus the bladders from mice receiving only ifosfamide) and several gamma interferon-inducible proteins. Also, the interferon signaling pathway was the most downregulated pathway in the bladders of IPSE-pretreated mice as compared to bladders only exposed to ifosfamide. Our results point to an association between downregulation of the interferon signaling pathway and amelioration of acrolein-induced urotoxicity, a mechanism that has not been previously linked to protection from hemorrhagic cystitis. The interferon signaling pathway has been previously shown to cross-talk with inflammasomes activated during inflammatory responses to irritants (95, 96). Some of these gamma interferon inducible genes are in turn linked to the development of pyroptosis, a highly inflammatory form of programmed cell death (Fig. 6B). Acrolein-induced pyroptotic cell death is a major determining factor of the severity of ifosfamide-induced urotoxicity (12, 32, 59). Pyroptosis can be compounded by activation of the inflammasome complex, which generates reactive species which perpetuate a vicious cycle of cell death (34, 36). The ability of IPSE to downregulate the interferon pathway, in conjunction with downregulation of major inflammatory pathways, may limit inflammasome activation and thus reduce acrolein-induced pyroptotic cell death. We hypothesize that IPSE, by bringing these processes to heel, subsequently modulates downstream urothelial damage, hemorrhage, oxidative stress and cellular infiltration.

The therapeutic efficacy of IPSE in ifosfamide-induced hemorrhagic cystitis may also partially depend on its modulation of oxidative stress cascades. Acrolein is a potent inducer of oxidative stress (33). Indeed, the NRF2-mediated oxidative stress responses pathway, which restores heme homeostasis and antioxidant responses, was highly expressed in the setting of ifosfamide-induced hemorrhagic cystitis (37). Accordingly, *Hmox1* and *Slc7a11* were among the top upregulated genes from this transcriptomic analysis. *Slc7a11* is a cysteine transporter, which has been implicated in glutathione metabolism in the bladder (80, 81). NRF2 induces the expression of heme oxygenase 1 (HO-1), the first enzyme of the heme oxygenase pathway, and several antioxidant enzymes including glutathione reductase (GSR), thioredoxin (TXN), thioredoxin reductase (TRXR1), superoxide dismutase (SOD), peroxiredoxin 1 (PRDX1), ferritin light chain (FTL) and ferritin heavy chains (FTH) (38). An association between an increase in the expression of NRF2 and protection from ifosfamide-induced hemorrhagic cystitis is consistent with a previous report linking hemostasis to reduction in hemorrhagic cystitis (37, 51, 53). In addition, there is a strong pathophysiological relationship between inflammation and oxidative stress (97). Severe pyroptosis can lead to enzymatic tissue damage and cellular DNA damage, which generate reactive species and superoxide radicals that induce oxidative stress (12, 97). When the bladder vasculature is exposed and injured following inflammation-driven urothelial damage, the resulting hemorrhage and release of heme further promotes oxidative stress. Based on the rationale that limiting inflammation reduces oxidative stress, and restoration of oxidative homeostasis initiates tissue repair processes, hemostatic agents and antioxidants have been widely tested as alternative therapies for preventing or reducing ifosfamide-induced hemorrhagic cystitis (37, 48-59). The link between inflammation and oxidative stress is evident from findings that most antioxidants showing efficacy in ifosfamide-induced hemorrhagic cystitis also downregulate pro-inflammatory cytokines and their downstream mediators (53, 56, 58, 61). In the same vein, anti-inflammatory drug candidates with efficacy in ifosfamide-induced hemorrhagic cystitis can also restore antioxidant enzyme activity to homoeostatic levels (43, 48, 49). In this study, however, we did not observe overt differential transcriptional changes in oxidative stress responses induced by IPSE pretreatment, although some antioxidant enzymes in bladders of IPSE-treated mice returned to basal levels. Nevertheless, it was interesting to observe that the genes encoding the proteins involved in iron homeostasis (*Fth* and *Ftl*) were relatively restored to baseline in bladders from the IPSE treated group. Also, some antioxidant proteins involved in xenobiotic detoxification and hemostasis (*Gclm, Gclc, Gsr* and *Gm3776*), superoxide detoxification (*Sod2*) and stress induced chaperones (*Stip1* and *Dnajb1*) were also reduced to normal levels in bladders from the IPSE-pretreated group. In addition, multidrug resistance protein 1 (*Abcc1, Mrp1*), which functions as an anion transporter with glutathione as a substrate (98), returned to basal levels in the ifosfamide-exposed bladder following IPSE pretreatment. Taken together, these differences suggest reduced levels of oxidative stress are present in ifosfamide-exposed bladders from IPSE-pretreated group, evident in reduced levels of genes related to detoxification of xenobiotics, DNA damage sensing and iron homeostasis.

Ifosfamide injury of the bladder may disrupt homeostasis of pathways besides oxidative stress. The PPAR pathway was downregulated in the ifosfamide-exposed bladder, which would presumably impair restoration of lipid homeostasis following epithelial membrane damage by acrolein. The PPAR pathway is also a modulator of inflammatory responses (86, 87) in addition to its role in the development and maintenance of IL-4 dependent alternatively activated status in macrophages (88), which we speculate may be inhibited in the highly inflammatory environment of the ifosfamide-injured bladder. Interestingly, we did not observe any significant effects of IPSE pretreatment on transcription of gene members of the PPAR pathway, suggesting a post-translational mechanism not captured by our early time point. In contrast, the most IPSE-downregulated genes in the ifosfamide-exposed bladder were those related to the neuro-inflammation signaling pathway (specifically, pathways active in central nervous system cells, i.e., astrocytes and microglia). We have previously reported that IPSE alleviates ifosfamide-induced allodynia (42). It remains to be shown whether this observed downregulation of neuro-inflammatory signaling is directly linked to this protective effect on allodynia, especially given the absence of astrocytes and microglia in the bladder.

While this study has revealed potential mechanistic changes associated with ifosfamide-induced hemorrhagic cystitis, and presented evidence of possible underlying mechanisms of IPSE’s therapeutic effect, our RNA-Seq-based approach cannot establish a causal relationship between observed gene expression and phenomena of interest (including IPSE’s therapeutic effects). This dataset did not reveal whether the therapeutic effects of IPSE are due to its IL-4-inducing properties, chemokine sequestration, or nuclear translocation-related, direct transcriptional effects. For instance, the downregulation of interferon signaling and its related genes is intriguing, but it is unclear to which extent this can be ascribed to IPSE-induced IL-4. Also, RNA-Seq does not capture epigenetic or post-translational regulation of gene or protein expression and activity. Moreover, we focused on a single, early time point following ifosfamide exposure. Although we observed differential transcription of multiple genes of interest at this time point, it is unlikely that this cross-sectional analysis has captured all relevant gene expression. Finally, this study focused on transcriptional changes in the bladder alone, and did not examine systemic gene expression induced by ifosfamide and IPSE. It is possible that such gene expression may account for some of ifosfamide and IPSE’s *in vivo* effects.

In conclusion, we have elucidated transcriptional dynamics associated with ifosfamide-induced hemorrhagic cystitis. These data provide new insights into the underlying mechanisms driving acrolein-induced urotoxicity associated with the use of ifosfamide and other oxazaphosphorines. We also showed that IPSE, an anti-inflammatory, parasite-derived molecule with therapeutic potential for ifosfamide-induced hemorrhagic cystitis (42), downregulates major inflammatory pathways potentially related to its mechanisms of effect. Our work demonstrates that there may be therapeutic potential for naturally occurring anti-inflammatory molecules, including pathogen-derived factors, as alternative or complementary therapies for ifosfamide-induced hemorrhagic cystitis. Apart from inhibition of inflammation and modest restoration of normal levels of antioxidants, we did not observe complete prevention of acrolein-induced oxidative stress by IPSE pretreatment. This is probably due to IPSE’s inability to directly bind to and neutralize acrolein (the mechanism of MESNA). Thus, IPSE is playing only a limited role on oxidative stress while suppressing inflammation. However, we have only compared one dose of IPSE given before ifosfamide challenge against three doses of MESNA. It remains to be shown whether IPSE will produce more ameliorative effects when given in multiple doses or through alternative routes. Ongoing work is focusing on optimization of IPSE, specifically related to its IL-4 induction and chemokine binding properties, to enhance its efficacy while preventing toxicity. Our hope is that these variations on IPSE and its administration will result in significantly improved efficacy, and ultimately, an alternative to MESNA in preventing ifosfamide-induced hemorrhagic cystitis.

## Materials and Methods

### Ethical Approval

Animal experiments reported in this study were conducted in a humane manner, adhering to relevant U.S. and international guidelines. Our animal handling and experimental protocols were reviewed and approved by the Institutional Animal Care and Use Committee (IACUC) of the Biomedical Research Institute, Rockville, Maryland, USA. Our IACUC guidelines comply with the U.S. Public Health Service Policy on Human Care and Use of Laboratory Animals.

### Animals, reagents and drugs

Female 7-week-old C57BL/6 mice (Charles River Laboratories, Wilmington, MA, USA) were housed using 12-h light-dark cycles in temperature-controlled holding rooms, with an unlimited supply of dry mouse chow and water. Ifosfamide (>98% purity) was purchased from Sigma-Aldrich (Sigma-Aldrich, St. Louis, MO, USA). IPSE cloning, expression and purification was performed as previously described (42, 78).

### Ifosfamide-induced hemorrhagic cystitis model

The ifosfamide-induced hemorrhagic cystitis model presented in this study was performed following methods previously described by Macedo et al., 2012 (25). Mice were intravenously injected with 25μg of IPSE or saline 24 hours before intraperitoneal ifosfamide injection (400mg/kg). Mice were then monitored for 6 hours post-ifosfamide injection before they were sacrificed for downstream experiments. Bladders were aseptically collected for RNA purification.

### RNA purification

RNA was isolated from mouse bladders using TRIzol Reagent and PureLink RNA Mini Kit (Invitrogen), according to manufacturers’ instructions. Briefly, aseptically excised bladders were homogenized in 1 ml TRIzol Reagent by bead-beating using ceramic beads (Omni International) and a mini-beadbeater (Biospec). Following a 5-min incubation, 0.2 ml chloroform was added and again incubated for 3 min before centrifugation at 12,000 ×g for 15 min to separate homogenates into aqueous and organic phases. The aqueous supernatant (~400ul) was mixed with an equal volume of 70% ethanol before binding the mixture to RNA binding columns by centrifugation. On-column DNase digestion (Invitrogen) was performed for 30 minutes, following the manufacturer’s protocols. After column washes and drying, RNA was eluted in RNase-free water, quantified and its quality checked using a NanoDrop 1000 spectrophotometer (Thermo Scientific) and Bioanalyzer 2100 (Agilent).

### RNA sequencing and RNA-seq analysis pipeline

RNA sequencing was performed using the Illumina-HiSeq2500/4000 NGS platform at a depth of ~20 million reads. Analyses were conducted using the RNA analysis tools of the Galaxy platform (www.usegalaxy.org). Raw sequence reads were aligned to the mouse genome (Mm10) by HISAT2 (99). The resulting alignment files, along with the corresponding mouse genome annotation file, were used as the input for HTSeq-count (100). DESeq2 (101) was used to determine differentially expressed genes between each pair of treatment groups. PCA plots were also generated by DESeq2. The DESeq2 results files containing gene IDs, log2 fold change and standard deviation, p-values and adjusted p-values were processed further downstream for functional analysis.

### Functional and pathway analysis, statistics and plots

Pathway, mechanistic network and functional analyses were generated using Ingenuity Pathways Analysis (QIAGEN Inc., https://www.qiagenbio-informatics.com/products/ingenuity-pathway-analysis) (102). The threshold cut-off was set at adjusted p-value < 0.1 for gene expression comparisons between bladders exposed to ifosfamide versus saline vehicle, and p < 0.05 for gene expression comparisons between IPSE-pretreated, ifosfamide-exposed bladders versus bladders only exposed to ifosfamide. The cut off for log2(fold change) was set at > 1 (2 fold). Other data analyses and plots were generated using GraphPad Prism v 6.00, and *ggplot2* and *plotly* packages in R. For comparisons among groups, one way analysis of variance (ANOVA) was performed and if significant, was followed by *post hoc* Student *t*-tests for pairwise comparisons after confirming a normal distribution. Plotted data show individual data points with error bars representing means and standard deviation.

### Histology

Bladders were fixed in 10% neutral-buffered formalin and later dehydrated and embedded in paraffin. Paraffin-embedded bladders were cut into 5 micron sections and then processed for hematoxylin and eosin staining. The stained sections were evaluated microscopically (in a blinded fashion by J.I.O.) for the presence of urothelial denudation, lamina propria edema, hemorrhage, and cellular infiltration.

## Acknowledgments

We gratefully acknowledge our funding sources, the Margaret A. Stirewalt Endowment (MHH), NIDDK R01DK113504 (MHH), NIAID R56AI119168 (MHH), and a Urology Care Foundation Research Scholar Award (ECM).

## Author contributions

Designed research studies (ECM, MHH), conducted experiments (ECM, LL, RZ, LFP, AA), acquired data (ECM, LL, RZ, KI, NB, LFP, AA, JIO), analyzed data (ECM, KI, NB, LL, LFP, MHH, JIO), providing reagents (LFP, TSJ, FHF, MHH), and wrote the manuscript (ECM, KI, TSJ, FHF, MHH).

## Supplementary Figure Legends

***Supplementary Fig. S1. Summary of the Pathways, Functional and Network analysis using Ingenuity Pathway Analysis.** This file shows top 5 each of canonical pathways, upstream regulators, diseases and disorders, molecular and cellular functions, physiological system development and functions, tox functions (hepatotoxicity, nephrotoxicity and cardiotoxicity), regulator effect networks, mechanistic networks, top 10 upregulated and downregulated genes*.

***Supplementary Fig. S2. Major upregulated pro-inflammatory pathways during ifosfamide induced hemorrhagic cystitis.** Following ifosfamide injection and acrolein induced urotoxicity, there was upregulation of upstream cytokines (IL-6, IL-1*β* and TNF*α*), their receptors, adaptor proteins, protein kinases and nuclear transcriptional factor (STAT3 and NFκB) in the (A) STAT3 pathway and (B) NFκB pathway. Keys: upregulation (red), downregulation (green), cytokines (square), growth factors (dotted square), phosphatase (triangle), kinases (inverted triangle), transmembrane receptors (ellipse), transcriptional regulators (wide circle), peptidase (rhombus), group or complex (double lined shapes), transporter (trapezium), acts on (line with filled arrow), translocate (line with open arrow), inhibition (line with perpendicular line at edge)*.

***Supplementary Fig. S3. Other upregulated pro-inflammatory pathways during ifosfamide induced hemorrhagic cystitis. Other major upregulated proinflammatory** pathways associated with ifosfamide induced hemorrhagic cystitis were Role of IL-17F in Allergic Airway Diseases, p38 MAPK signaling, Leucocyte Extravasation signaling, HMGB1 signaling, TREM1 signaling. For the key to the annotations, see description in Supplementary Fig. S2 legend*.

***Supplementary Fig. S4. NRF2 mediated oxidative stress responses pathway.** NRF2 is the major pathway regulating response to oxidative stress. It induces the expression of heme oxygenase pathway, the first enzyme of the heme homoeostasis pathway, and the expression of several antioxidant enzymes and proteins. An abridged version of this figure is shown in Fig. 5. For the key to the annotations, see description in Supplementary Fig. S2 legend*.

***Supplementary Fig. S5. PPAR signaling pathway.** This is the major pathway regulating lipid homoeostasis. PPAR has been shown to play an anti-inflammatory role (87), thus, here downregulated in response to ifosfamide induced cystitis. For the key to the annotations, see description in Supplementary Fig. S2 legend*.

***Supplementary Fig. S6. Diseases and Function Tree map.** This is a graphical representation of changes in the diseases and disorders, molecular and cellular functions, physiological system development and functions altered due to ifosfamide induced cystitis. We saw high upregulation of functions related to organismal injury and abnormalities, inflammatory diseases, cancer, cell proliferation, cellular movement and hematological systems development and function, and downregulation of cell death in response to ifosfamide induced cystitis*.

***Supplementary Fig. S7. Neuro-inflammation pathway.** There was potential neurotoxic effect due to ifosfamide induced hemorrhagic cystitis. We observed notable upregulation of proinflammatory cytokines and pro-inflammatory mediators in astrocytes and microglia cells. This is consistent with the known neurotoxic effect of ifosfamide. For the key to the annotations, see description in Supplementary Fig. S2 legend*.

***Supplementary Fig. S8. Hypoxia signaling pathway.** As a result of hemorrhage, there was indication of hypoxia in the bladder as depicted by the upregulation of HIF-1*α* mediated hypoxia. For the key to the annotations, see description in Supplementary Fig. S2 legend*.

***Supplementary Fig. S9. Mechanistic network analysis.** Mechanistic network analysis of transcriptome of IPSE pretreated mice compared to ifosfamide only mice showed downregulation of interactions between several proinflammatory genes. In addition to the network interaction between chemokines and interferon induced proteins, we also recorded more downregulatory mechanistic network interaction between genes encoding interferons induced proteins. For the key to the annotations, see description in Supplementary Fig. S2 legend*.

## Abbreviations

IPSE: interleukin-4 inducing principle from *Schistosoma* eggs
MESNA: 2-mercaptoethane sulfonate Sodium

## References

1. Young SD, Whissell M, Noble JCS, Cano PO, Lopez PG, and Germond CJ. Phase II clinical trial results involving treatment with low-dose daily oral cyclophosphamide, weekly vinblastine, and rofecoxib in patients with advanced solid tumors. Clinical Cancer Research. 2006;12:3092–8.

2. Advani SH. The role of ifosfamide in paediatric cancer. Aust N Z J Med. 1998;28(3):410–3.

3. Lawson M, Vasilaras A, De Vries A, Mactaggart P, and Nicol D. Urological implications of cyclophosphamide and ifosfamide. Scandinavian Journal of Urology and Nephrology. 2008;42(4):309–17.

4. Traxer O, Desgrandchamps F, Sebe P, Haab F, Le Duc A, Gattegno B, et al. [Hemorrhagic cystitis: etiology and treatment]. Prog Urol. 2001;11(4):591–601.

5. Mendenhall WM, Henderson RH, Costa JA, Hoppe BS, Dagan R, Bryant CM, et al. Hemorrhagic radiation cystitis. Am J Clin Oncol. 2015;38(3):331–6.

6. Rajaganapathy BR, Jayabalan N, Tyagi P, Kaufman J, and Chancellor MB. Advances in Therapeutic Development for Radiation Cystitis. Low Urin Tract Symptoms. 2014;6(1):1–10.

7. Alesawi AM, El-Hakim A, Zorn KC, and Saad F. Radiation-induced hemorrhagic cystitis. Curr Opin Support Palliat Care. 2014;8(3):235–40.

8. Arora R, Jasmita Singh M, Garg A, Gupta M, and Gupta N. Successful Treatment of BK Virus Hemorrhagic Cystitis (HC) Post Allogenic Hematopoietic Stem Cell Transplantation with Low Dose Cidofovir. J Assoc Physicians India. 2017;65(5):93–4.

9. Kato J, Mori T, Suzuki T, Ito M, Li TC, Sakurai M, et al. Nosocomial BK Polyomavirus Infection Causing Hemorrhagic Cystitis Among Patients With Hematological Malignancies After Hematopoietic Stem Cell Transplantation. Am J Transplant. 2017;17(9):2428–33.

10. Perez-Huertas P, Cueto-Sola M, Escobar-Cava P, Fernandez-Navarro JM, Borrell-Garcia C, Albert-Mari A, et al. BK Virus-Associated Hemorrhagic Cystitis After Allogeneic Hematopoietic Stem Cell Transplantation in the Pediatric Population. J Pediatr Oncol Nurs. 2016.

11. delaCruz J, and Pursell K. BK Virus and Its Role in Hematopoietic Stem Cell Transplantation: Evolution of a Pathogen. Curr Infect Dis Rep. 2014;16(8):417.

12. Haldar S, Dru C, and Bhowmick NA. Mechanisms of hemorrhagic cystitis. Am J Clin Exp Urol. 2014;2(3):199–208.

13. Sarosy G. Ifosfamide--pharmacologic overview. Semin Oncol. 1989;16(1 Suppl 3):2–8.

14. Sakurai M, Saijo N, Shinkai T, Eguchi K, Sasaki Y, Tamura T, et al. The protective effect of 2-mercapto-ethane sulfonate (MESNA) on hemorrhagic cystitis induced by high-dose ifosfamide treatment tested by a randomized crossover trial. Jpn J Clin Oncol. 1986;16(2):153–6.

15. Andriole GL, Sandlund JT, Miser JS, Arasi V, Linehan M, and Magrath IT. The efficacy of mesna (2-mercaptoethane sodium sulfonate) as a uroprotectant in patients with hemorrhagic cystitis receiving further oxazaphosphorine chemotherapy. Journal of clinical oncology: official journal of the American Society of Clinical Oncology. 1987;5(5):799–803.

16. Higgs D, Nagy C, and Einhorn LH. Ifosfamide: a clinical review. Semin Oncol Nurs. 1989;5(2 Suppl 1):70–7.

17. Shepherd JD, Pringle LE, Barnett MJ, Klingemann H-G, Reece DE, and Phillips GL. Mesna versus hyperhydration for the prevention of cyclophosphamide- induced hemorrhagic cystitis in bone marrow transplantation. Journal of Clinical Oncology. 1991;9:2016–20.

18. Furlanut M, and Franceschi L. Pharmacology of ifosfamide. Oncology. 2003;65 Suppl 2:2–6.

19. Shimogori K, Araki M, Shibazaki S, Tuda K, and Miura K. Nonimmediate allergic reactions induced by Mesna. J Gen Fam Med. 2017;18(5):285–7.

20. Behnam K, Patil UB, and Mariano E. Intravesical instillation of Formalin for hemorrhagic cystitis secondary to radiation for gynecologic malignancies. Gynecol Oncol. 1983;16(1):31–3.

21. Donahue LA, and Frank IN. Intravesical formalin for hemorrhagic cystitis: analysis of therapy. J Urol. 1989;141(4):809–12.

22. Montgomery BD, Boorjian SA, Ziegelmann MJ, Joyce DD, and Linder BJ. Intravesical silver nitrate for refractory hemorrhagic cystitis. Turk J Urol. 2016;42(3):197–201.

23. Ziegelmann MJ, Boorjian SA, Joyce DD, Montgomery BD, and Linder BJ. Intravesical formalin for hemorrhagic cystitis: A contemporary cohort. Can Urol Assoc J. 2017;11(3-4):E79–e82.

24. Russo P. Urologic emergencies in the cancer patient. Semin Oncol. 2000;27(3):284–98.

25. Macedo FY, Mourao LT, Freitas HC, Lima RC, Jr., Wong DV, Oria RB, et al. Interleukin-4 modulates the inflammatory response in ifosfamide-induced hemorrhagic cystitis. Inflammation. 2012;35(1):297–307.

26. Levenback C, Eifel PJ, Burke TW, Morris M, and Gershenson DM. Hemorrhagic cystitis following radiotherapy for stage Ib cancer of the cervix. Gynecol Oncol. 1994;55(2):206–10.

27. Kaplan JR, and Wolf JS, Jr. Efficacy and survival associated with cystoscopy and clot evacuation for radiation or cyclophosphamide induced hemorrhagic cystitis. J Urol. 2009;181(2):641–6.

28. Kaur D, Khan SP, Rodriguez V, Arndt C, and Claus P. Hyperbaric oxygen as a treatment modality in cyclophosphamide-induced hemorrhagic cystitis. Pediatr Transplant. 2018:e13171.

29. Andriole GL, Yuan JJ, and Catalona WJ. Cystotomy, temporary urinary diversion and bladder packing in the management of severe cyclophosphamide-induced hemorrhagic cystitis. J Urol. 1990;143(5):1006–7.

30. West NJ. Prevention and treatment of hemorrhagic cystitis. Pharmacotherapy. 1997;17(4):696–706.

31. Matz EL, and Hsieh MH. Review of Advances in Uroprotective Agents for Cyclophosphamide- and Ifosfamide-induced Hemorrhagic Cystitis. Urology. 2017;100:16–9.

32. Korkmaz A, Topal T, and Oter S. Pathophysiological aspects of cyclophosphamide and ifosfamide induced hemorrhagic cystitis; Implication of reactive oxygen and nitrogen species as well as PARP activation. Cell Biology and Toxicology. 2007;23:303–12.

33. Moghe A, Ghare S, Lamoreau B, Mohammad M, Barve S, McClain C, et al. Molecular mechanisms of acrolein toxicity: relevance to human disease. Toxicol Sci. 2015;143(2):242–55.

34. Poeck H, Bscheider M, Gross O, Finger K, Roth S, Rebsamen M, et al. Recognition of RNA virus by RIG-I results in activation of CARD9 and inflammasome signaling for interleukin 1 beta production. Nat Immunol. 2010;11(1):63–9.

35. Gomes TN, Santos CC, Souza-Filho MV, Cunha FQ, and Ribeiro RA. Participation of TNF-alpha and IL-1 in the pathogenesis of cyclophosphamide-induced hemorrhagic cystitis. Braz J Med Biol Res. 1995;28(10):1103–8.

36. Takeuchi O, and Akira S. Pattern recognition receptors and inflammation. Cell. 2010;140(6):805–20.

37. Gore PR, Prajapati CP, Mahajan UB, Goyal SN, Belemkar S, Ojha S, et al. Protective Effect of Thymoquinone against Cyclophosphamide-Induced Hemorrhagic Cystitis through Inhibiting DNA Damage and Upregulation of Nrf2 Expression. Int J Biol Sci. 2016;12(8):944–53.

38. Nguyen T, Nioi P, and Pickett CB. The Nrf2-antioxidant response element signaling pathway and its activation by oxidative stress. J Biol Chem. 2009;284(20):13291–5.

39. Hingorani P, Zhang W, Piperdi S, Pressman L, Lin J, Gorlick R, et al. Preclinical activity of palifosfamide lysine (ZIO-201) in pediatric sarcomas including oxazaphosphorine-resistant osteosarcoma. Cancer Chemother Pharmacol. 2009;64(4):733–40.

40. Kopp HG, Kanz L, and Hartmann JT. Hypersensitivity pneumonitis associated with the use of trofosfamide. Anticancer Drugs. 2004;15(6):603–4.

41. Leite CA, Alencar VT, Melo DL, Mota JM, Melo PH, Mourao LT, et al. Target Inhibition of IL-1 Receptor Prevents Ifosfamide Induced Hemorrhagic Cystitis in Mice. J Urol. 2015;194(6):1777–86.

42. Mbanefo EC, Le L, Pennington LF, Odegaard JI, Jardetzky TS, Alouffi A, et al. Therapeutic exploitation of IPSE, a urogenital parasite-derived host modulatory protein, for chemotherapy-induced hemorrhagic cystitis. Faseb j. 2018:fj201701415R.

43. Kim SH, Lee IC, Ko JW, Moon C, Kim SH, Shin IS, et al. Diallyl Disulfide Prevents Cyclophosphamide-Induced Hemorrhagic Cystitis in Rats through the Inhibition of Oxidative Damage, MAPKs, and NF-kappaB Pathways. Biomol Ther (Seoul). 2015;23(2):180–8.

44. Wrobel A, Doboszewska U, Rechberger E, Rojek K, Serefko A, Poleszak E, et al. Rho kinase inhibition ameliorates cyclophosphamide-induced cystitis in rats. Naunyn Schmiedebergs Arch Pharmacol. 2017;390(6):613–9.

45. Dietrich F, Pietrobon Martins J, Kaiser S, Madeira Silva RB, Rockenbach L, Albano Edelweiss MI, et al. The Quinovic Acid Glycosides Purified Fraction from Uncaria tomentosa Protects against Hemorrhagic Cystitis Induced by Cyclophosphamide in Mice. PLoS One. 2015;10(7):e0131882.

46. Santos AA, Jr., Leal PC, Edelweiss MI, Lopes TG, Calixto JB, Morrone FB, et al. Effects of the compounds MV8608 and MV8612 obtained from Mandevilla velutina in the model of hemorrhagic cystitis induced by cyclophosphamide in rats. Naunyn Schmiedebergs Arch Pharmacol. 2010;382(5-6):399–407.

47. Dantas AC, Batista-Junior FF, Macedo LF, Mendes MN, Azevedo IM, and Medeiros AC. Protective effect of simvastatin in the cyclophosphamide-induced hemorrhagic cystitis in rats. Acta Cir Bras. 2010;25(1):43–6.

48. Sherif IO, Nakshabandi ZM, Mohamed MA, and Sarhan OM. Uroprotective effect of oleuropein in a rat model of hemorrhagic cystitis. Int J Biochem Cell Biol. 2016;74:12–7.

49. Abo-Salem OM. Uroprotective effect of pentoxifylline in cyclophosphamide-induced hemorrhagic cystitis in rats. J Biochem Mol Toxicol. 2013;27(7):343–50.

50. Kilic O, Akand M, Karabagli P, and Piskin MM. Hemostatic Efficacy and Histopathological Effects of Ankaferd Blood Stopper in an Experimental Rat Model of Cyclophosphamide-induced Hemorrhagic Cystitis. Urology. 2016;94:313.e7-.e13.

51. Chow YC, Yang S, Huang CJ, Tzen CY, Huang PL, Su YH, et al. Epinephrine promotes hemostasis in rats with cyclophosphamide-induced hemorrhagic cystitis. Urology. 2006;67(3):636–41.

52. Chow YC, Yang S, Huang CJ, Tzen CY, Su YH, and Wang PS. Prophylactic intravesical instillation of epinephrine prevents cyclophosphamide-induced hemorrhagic cystitis in rats. Exp Biol Med (Maywood). 2007;232(4):565–70.

53. Matsuoka Y, Masuda H, Yokoyama M, and Kihara K. Protective effects of bilirubin against cyclophosphamide induced hemorrhagic cystitis in rats. J Urol. 2008;179(3):1160–6.

54. Kanat O, Kurt E, Yalcinkaya U, Evrensel T, and Manavoglu O. Comparison of uroprotective efficacy of mesna and amifostine in Cyclophosphamide- induced hemorrhagic cystitis in rats. Indian J Cancer. 2006;43(1):12–5.

55. Boeira VT, Leite CE, Santos AA, Jr., Edelweiss MI, Calixto JB, Campos MM, et al. Effects of the hydroalcoholic extract of Phyllanthus niruri and its isolated compounds on cyclophosphamide-induced hemorrhagic cystitis in mouse. Naunyn Schmiedebergs Arch Pharmacol. 2011;384(3):265–75.

56. Hamsa TP, and Kuttan G. Protective role of Ipomoea obscura (L.) on cyclophosphamide-induced uro- and nephrotoxicities by modulating antioxidant status and pro-inflammatory cytokine levels. Inflammopharmacology. 2011;19(3):155–67.

57. Yildirim I, Korkmaz A, Oter S, Ozcan A, and Oztas E. Contribution of antioxidants to preventive effect of mesna in cyclophosphamide-induced hemorrhagic cystitis in rats. Cancer Chemother Pharmacol. 2004;54(5):469–73.

58. Sadir S, Deveci S, Korkmaz A, and Oter S. Alpha-tocopherol, beta-carotene and melatonin administration protects cyclophosphamide-induced oxidative damage to bladder tissue in rats. Cell Biochem Funct. 2007;25(5):521–6.

59. Zupancic D, Jezernik K, and Vidmar G. Effect of melatonin on apoptosis, proliferation and differentiation of urothelial cells after cyclophosphamide treatment. J Pineal Res. 2008;44(3):299–306.

60. Ozguven AA, Yilmaz O, Taneli F, Ulman C, Vatansever S, and Onag A. Protective effect of ketamine against hemorrhagic cystitis in rats receiving ifosfamide. Indian J Pharmacol. 2014;46(2):147–51.

61. Sakura M, Masuda H, Matsuoka Y, Yokoyama M, Kawakami S, and Kihara K. Rolipram, a specific type-4 phosphodiesterase inhibitor, inhibits cyclophosphamide-induced haemorrhagic cystitis in rats. BJU Int. 2009;103(2):264–9.

62. Yigitaslan S, Ozatik O, Ozatik FY, Erol K, Sirmagul B, and Baseskioglu AB. Effects of tadalafil on hemorrhagic cystitis and testicular dysfunction induced by cyclophosphamide in rats. Urol Int. 2014;93(1):55–62.

63. Vela-Ojeda J, Tripp-Villanueva F, Sanchez-Cortes E, Ayala-Sanchez M, Garcia-Ruiz Esparza MA, Rosas-Cabral A, et al. Intravesical rhGM-CSF for the treatment of late onset hemorrhagic cystitis after bone marrow transplant. Bone Marrow Transplant. 1999;24(12):1307–10.

64. Mota JM, Brito GA, Loiola RT, Cunha FQ, and Ribeiro RdA. Interleukin-11 attenuates ifosfamide-induced hemorrhagic cystitis. International braz j urol: official journal of the Brazilian Society of Urology. 33:704–10.

65. Ozyuvali E, Yildirim ME, Yaman T, Kosem B, Atli O, and Cimentepe E. Protective Effect of Intravesical Platelet-Rich Plasma on Cyclophosphamide-Induced Hemorrhagic Cystitis. Clin Invest Med. 2016;39(6):27514.

66. Donmez MI, Inci K, Zeybek ND, Dogan HS, and Ergen A. The Early Histological Effects of Intravesical Instillation of Platelet-Rich Plasma in Cystitis Models. Int Neurourol J. 2016;20(3):188–96.

67. Freitas RD, Costa KM, Nicoletti NF, Kist LW, Bogo MR, and Campos MM. Omega-3 fatty acids are able to modulate the painful symptoms associated to cyclophosphamide-induced-hemorrhagic cystitis in mice. J Nutr Biochem. 2016;27:219–32.

68. Keles I, Bozkurt MF, Cemek M, Karalar M, Hazini A, Alpdagtas S, et al. Prevention of cyclophosphamide-induced hemorrhagic cystitis by resveratrol: a comparative experimental study with mesna. Int Urol Nephrol. 2014;46(12):2301–10.

69. Vieira MM, Macedo FY, Filho JN, Costa AC, Cunha AN, Silveira ER, et al. Ternatin, a flavonoid, prevents cyclophosphamide and ifosfamide-induced hemorrhagic cystitis in rats. Phytother Res. 2004;18(2):135–41.

70. Assreuy AM, Martins GJ, Moreira ME, Brito GA, Cavada BS, Ribeiro RA, et al. Prevention of cyclophosphamide-induced hemorrhagic cystitis by glucose-mannose binding plant lectins. J Urol. 1999;161(6):1988–93.

71. Arafa HM. Uroprotective effects of curcumin in cyclophosphamide-induced haemorrhagic cystitis paradigm. Basic Clin Pharmacol Toxicol. 2009;104(5):393–9.

72. Meyer NH, Mayerhofer H, Tripsianes K, Blindow S, Barths D, Mewes A, et al. A Crystallin Fold in the Interleukin-4- inducing Principle of Schistosoma mansoni Eggs (IPSE/alpha-1) Mediates IgE Binding for Antigen-independent Basophil Activation. The Journal of biological chemistry. 2015;290(36):22111–26.

73. Schramm G, Falcone FH, Gronow A, Haisch K, Mamat U, Doenhoff MJ, et al. Molecular characterization of an interleukin- 4-inducing factor from Schistosoma mansoni eggs. The Journal of biological chemistry. 2003;278(20):18384–92.

74. Schramm G, Mohrs K, Wodrich M, Doenhoff MJ, Pearce EJ, Haas H, et al. Cutting edge: IPSE/alpha-1, a glycoprotein from Schistosoma mansoni eggs, induces IgE-dependent, antigen-independent IL-4 production by murine basophils in vivo. Journal of immunology (Baltimore, Md: 1950). 2007;178(10):6023–7.

75. Smith P, Fallon RE, Mangan NE, Walsh CM, Saraiva M, Sayers JR, et al. Schistosoma mansoni secretes a chemokine binding protein with antiinflammatory activity. The Journal of experimental medicine. 2005;202(10):1319–25.

76. Kaur I, Schramm G, Everts B, Scholzen T, Kindle KB, Beetz C, et al. Interleukin-4-inducing principle from Schistosoma mansoni eggs contains a functional C-terminal nuclear localization signal necessary for nuclear translocation in mammalian cells but not for its uptake. Infection and immunity. 2011;79(4):1779–88.

77. Meyer NH, Schramm G, and Sattler M. 1H, 13C and 15N chemical shift assignments of IPSEDeltaNLS. Biomolecular NMR assignments. 2011;5(2):225–7.

78. Pennington LF, Alouffi A, Mbanefo EC, Ray D, Heery DM, Jardetzky TS, et al. H-IPSE is a pathogen-secreted host nucleus infiltrating protein (infiltrin) expressed exclusively by the Schistosoma haematobium egg stage. Infect Immun. 2017;85(12):e00301-17.

79. Magrini E, Mantovani A, and Garlanda C. The Dual Complexity of PTX3 in Health and Disease: A Balancing Act? Trends Mol Med. 2016;22(6):497–510.

80. Drayton RM, Dudziec E, Peter S, Bertz S, Hartmann A, Bryant HE, et al. Reduced expression of miRNA-27a modulates cisplatin resistance in bladder cancer by targeting the cystine/glutamate exchanger SLC7A11. Clin Cancer Res. 2014;20(7):1990–2000.

81. Wei L, Chintala S, Ciamporcero E, Ramakrishnan S, Elbanna M, Wang J, et al. Genomic profiling is predictive of response to cisplatin treatment but not to PI3K inhibition in bladder cancer patient-derived xenografts. Oncotarget. 2016;7(47):76374-89.

82. Dejima T, Shibata K, Yamada H, Takeuchi A, Hara H, Eto M, et al. A C-type lectin receptor pathway is responsible for the pathogenesis of acute cyclophosphamide-induced cystitis in mice. Microbiol Immunol. 2013;57(12):833–41.

83. Ribeiro RA, Freitas HC, Campos MC, Santos CC, Figueiredo FC, Brito GA, et al. Tumor necrosis factor-alpha and interleukin-1beta mediate the production of nitric oxide involved in the pathogenesis of ifosfamide induced hemorrhagic cystitis in mice. J Urol. 2002;167(5):2229–34.

84. Wang CC, Weng TI, Wu ET, Wu MH, Yang RS, and Liu SH. Involvement of interleukin-6-regulated nitric oxide synthase in hemorrhagic cystitis and impaired bladder contractions in young rats induced by acrolein, a urinary metabolite of cyclophosphamide. Toxicol Sci. 2013;131(1):302–10.

85. Dunning KR, Anastasi MR, Zhang VJ, Russell DL, and Robker RL. Regulation of fatty acid oxidation in mouse cumulus-oocyte complexes during maturation and modulation by PPAR agonists. PLoS One. 2014;9(2):e87327.

86. Reynders V, Loitsch S, Steinhauer C, Wagner T, Steinhilber D, and Bargon J. Peroxisome proliferator-activated receptor alpha (PPAR alpha) down-regulation in cystic fibrosis lymphocytes. Respir Res. 2006;7:104.

87. Cardell LO, Hagge M, Uddman R, and Adner M. Downregulation of peroxisome proliferator-activated receptors (PPARs) in nasal polyposis. Respir Res. 2005;6:132.

88. Chawla A. Control of macrophage activation and function by PPARs. Circ Res. 2010;106(10):1559–69.

89. Choi SH, Byun Y, and Lee G. Expressions of Uroplakins in the Mouse Urinary Bladder with Cyclophosphamide-Induced Cystitis. Journal of Korean Medical Science. 2009;24:684–9.

90. Lee G. Uroplakins in the lower urinary tract. Int Neurourol J. 2011;15:4–12.

91. Macedo FYB, Baltazar Ft, Almeida PRC, T??vora Fb, Ferreira FV, Schmitt FC, et al. Cyclooxygenase-2 expression on ifosfamide-induced hemorrhagic cystitis in rats. Journal of Cancer Research and Clinical Oncology. 2008;134:19–27.

92. Oter S, Korkmaz A, Oztas E, Yildirim I, Topal T, and Bilgic H. Inducible nitric oxide synthase inhibition in cyclophosphamide induced hemorrhagic cystitis in rats. Urol Res. 2004;32(3):185–9.

93. Sakthivel SK, Singh UP, Singh S, Taub DD, Novakovic KR, and Lillard JW, Jr. CXCL10 blockade protects mice from cyclophosphamide-induced cystitis. J Immune Based Ther Vaccines. 2008;6:6.

94. Dornelles FN, Andrade EL, Campos MM, and Calixto JB. Role of CXCR2 and TRPV1 in functional, inflammatory and behavioural changes in the rat model of cyclophosphamide-induced haemorrhagic cystitis. Br J Pharmacol. 2014;171(2):452–67.

95. Kopitar-Jerala N. The Role of Interferons in Inflammation and Inflammasome Activation. Front Immunol. 2017;8:873.

96. Malireddi RK, and Kanneganti TD. Role of type I interferons in inflammasome activation, cell death, and disease during microbial infection. Front Cell Infect Microbiol. 2013;3:77.

97. Biswas SK. Does the Interdependence between Oxidative Stress and Inflammation Explain the Antioxidant Paradox? Oxidative Medicine and Cellular Longevity. 2016;2016:9.

98. Cole SP. Multidrug resistance protein 1 (MRP1, ABCC1), a "multitasking" ATP-binding cassette (ABC) transporter. J Biol Chem. 2014;289(45):30880–8.

99. Kim D, Langmead B, and Salzberg SL. HISAT: a fast spliced aligner with low memory requirements. Nat Methods. 2015;12(4):357–60.

100. Anders S, Pyl PT, and Huber W. HTSeq--a Python framework to work with high-throughput sequencing data. Bioinformatics. 2015;31(2):166–9.

101. Love MI, Huber W, and Anders S. Moderated estimation of fold change and dispersion for RNA-seq data with DESeq2. Genome Biol. 2014;15(12):550.

102. Kramer A, Green J, Pollard J, Jr., and Tugendreich S. Causal analysis approaches in Ingenuity Pathway Analysis. Bioinformatics. 2014;30(4):523–30.

